# Sprr1 and miR-130b contribute to the senescence-like phenotype in aging

**DOI:** 10.1101/2023.10.25.563779

**Authors:** Ji Yeon Hong, Hee Jin Nam, Hong Ji, Yoon Young Kim, Miri Hyun, Hee Jung Park, Seung Min Bae, Pierre D. McCrea

**Author notes:** Co-corresponding authors during submission., Ji Yeon Hong, Ph.D. is the corresponding author upon publication: Pierre D. McCrea, Ph.D., Ji Yeon Hong, Ph.D. Daewoong Pharmaceutical, 72 Dugye-ro, Pogok-eup, Cheoin-gu, Yonginsi, Gyeonggi-do, 17028, Korea (Ji Yeon Hong).

## Abstract

Aging is an inevitable process with senescence being one of its hallmarks. Recent advances have indicated that the elimination of senescent cells can reduce the signs of aging and increase healthy life span. Here, we identify a negative modulator of aging, Sprr1a, and in turn a negative modulator of Sprr1a, miR-130b. We show that reductions in Sprr1a levels, including *via* miR-130b expression, promotes cell senescence-like phenotype. We find that mediators of senescence, such as inflammatory cytokines and cell cycle regulators, are modulated by the miR-130b and Sprr1a-related pathway. For example, the levels of p16, p53 and p21 become decreased or increased upon the respective expression of Sprr1a versus miR-130b. Further, as shown in relation to p16 levels and β-galactosidase levels, cells expressing Sprr1a exhibit significant protection from senescence-inducing factors such as radiation or Doxorubicin, suggesting that Sprr1a might contribute to protection against age-related pathologies. Taken together, we introduce two modulators of properties associated with senescence-like phenotype.

## INTRODUCTION

Aging is the major risk factor for a variety of adverse health outcomes such as cancer, cardiovascular disease, diabetes and neurodegenerative disorders [1–8]. Since aging populations are growing worldwide, it is relevant to identify proteins and pathways that contribute positively or negatively to aging, as well as linked phenotypes and biomarkers.

Cellular senescence was first identified by Hayflick [9, 10] during the serial passage of primary cells. Characteristics of senescence include a large and flat morphology and high acidic β-galactosidase enzyme activity accompanied by phenotypic alterations [11], such as chromatin remodeling, metabolic reprogramming, autophagy, and a proinflammatory secretome [12–16]. Cellular senescence is one of the hallmarks of aging [17–20]. Research has revealed that senescent cells accumulate in the mitotic tissues of aging primates [19–21], with new evidence indicating that the removal of senescent cells increases lifespan [22, 23]. This has encouraged experimentation with therapeutic approaches such as senolytics [24].

Senescent cells are metabolically active and secrete proinflammatory cytokines, chemokines and growth factors. Such signals released from senescent cells produce what is termed the <u>S</u>enescence <u>A</u>ssociated <u>S</u>ecretory <u>P</u>henotype (SASP) [16, 25]. Through autocrine or paracrine mechanisms, accumulated effects of SASP include the promotion and reinforcement of multiple senescence related properties. For example, contributing to aging phenotypes, SASP is thought to be partially responsible for persistent chronic inflammation known as inflammaging [26, 27], where IL-6 and IL-8 are among SASP effectors [28, 29].

Stable growth arrest is one of the hallmarks of senescence, with tumor suppressor pathways likely involved, namely p53 [28, 30], p21 [31, 32] and p16/Rb [33–36]. Intriguingly, the clearance of p16-positive senescent cells delays the onset of age-related pathologies in tissues such as adipose, skeletal muscle and eye [23]. In addition, by blocking the binding of FOXO4 to p53, apoptosis is promoted in senescent cells with the restoration of fitness, hair density and renal function in aged mice [22]. Such findings imply that the elimination of senescent cells aids in furthering healthy life span.

To address molecular mechanisms of aging, we compared gene and microRNA expression profiles of mesenchymal stromal cells (MSCs) isolated from young (4-6 months) *versus* aged mice (18-26 months). In our microarray data from older mice, we paid attention to down regulated gene transcripts as well as up regulated microRNAs. We focused upon the reduced expression of Sprr1a in MSCs from aged mice, in combination with increased expression of its potential negative modulator, miR-130b. With our findings including the capacity of Sprr1a to rescue cells from senescent-like phenotypes, we propose a model whereby the miR130b/ Sprr1a pathway modulates cellular senescence-like phenotype.

Sprr1a, also known as Cornifin A, is a small (∼15.8kDa mouse; ∼9.9kDa human) protein encoded by a regeneration-associated gene [37]. Its central domain contains proline-rich peptide repeats, while its amino- and carboxyl-terminal regions are enriched in glutamine and lysine. *In vitro*, Sprr1a promotes neural growth and it positively modulates nerve regeneration in conjunction with Sox11 [38]. To examine potential effectors of cellular senescence arising from our microarray findings, we focused upon Sprr1a. Summarized here but detailed below, our expression of Sprr1a decreased that of p53 and p16, in keeping with observed decreases in the SASP secretome. Conversely, the depletion of Sprr1a increased IL-6 levels, halting cell growth. Based upon homology pairing with the 3’ UTR of mouse Sprr1a, our *in silico* analysis (MicroCosm) initially left open the possibility that miR-130b might be a direct negative modulator of Sprr1a. However, experimental evidence, including the lack of cross-species conservation of such a site within the 3’ UTR, leads us to propose an indirect regulatory mechanism. miR-130b expression led to decreased levels of Sprr1a and increased levels of p16, p53, and the senescence marker, β-galactosidase. Furthermore, expression of miR-130b antagomir reduced the levels of senescence-related factors, as did the expression of Sprr1a. Finally, Sprr1a expression diminished cell responses to radiation and Doxorubicin. Together, we propose the existence of a miR-130b ➔ Sprr1a regulatory relationship that is likely indirect and contributes to cellular senescence-related phenotype regulation. We speculate in turn that this relationship or pathway responds to intracellular or extracellular age-associated stimuli.

## RESULTS

### Sprr1a expression inhibits senescence-associated phenotypes

Among the microarray candidate genes we resolved during the ageing of mouse mesenchymal stromal cells (MSCs), we concentrated upon Sprr1a (<u>S</u>mall <u>Pr</u>oline <u>R</u>ich protein 1A). This was due in part to Sprr1a’s considerably lower (9.8-fold) expression in aging, in combination with heightened expression of Sprr1a being previously associated with regeneration [39]. An additional aspect of intrigue is that Sprr1a’s downstream mechanism of action continues to be unknown. To evaluate whether Sprr1a modulates cell senescence, mouse Sprr1a was ectopically expressed in mouse MSCs isolated from aged mice (18-20 months). Generally, senescent cells feature an enlarged flattened cytoplasm and an irregular shape. Aged MSCs similarly exhibit such morphologies *in vitro* compared to MSCs isolated from young mice (2-3 months) (Supplemental Fig 1A). Surprisingly, exogenous expression of Sprr1a changed the morphology of old MSCs towards spindle shapes similar to those of MSCs from young mice (Fig 1A). In addition, cells which overexpress Sprr1a showed reduced β-galactosidase staining, one of the major biomarkers employed in detecting senescent cells [40]. We next evaluated the expression levels of select cell cycle proteins. P16 is known to be up-regulated when cells are senescent, and it is used as a biomarker of aging. The clearance of p16-positive cells extends medical lifespan, delays tumorigenesis and attenuates ageing [23]. In keeping with our initial data (Fig 1A), ectopic expression of Sprr1a reduced p16 protein levels (Fig 1B), consistent with the possibility that Sprr1a modulates cellular senescence. In addition, Sprr1a expression decreased the transcript levels of the cell-cycle related gene product p21 (Fig 1C). As described previously, one of the hallmarks of senescence cells is a senescence-associated secretory phenotype (SASP). Senescent cells secrete inflammatory cytokines such as IL-6, IL-8 and IL-1a, promoting age-associated inflammation and pathology. We found that the expression of Sprr1a in mouse MSCs decreased the mRNA levels of IL-6, and to a lesser but still notable extent IL-1L (Fig 1C).

**Figure 1.**
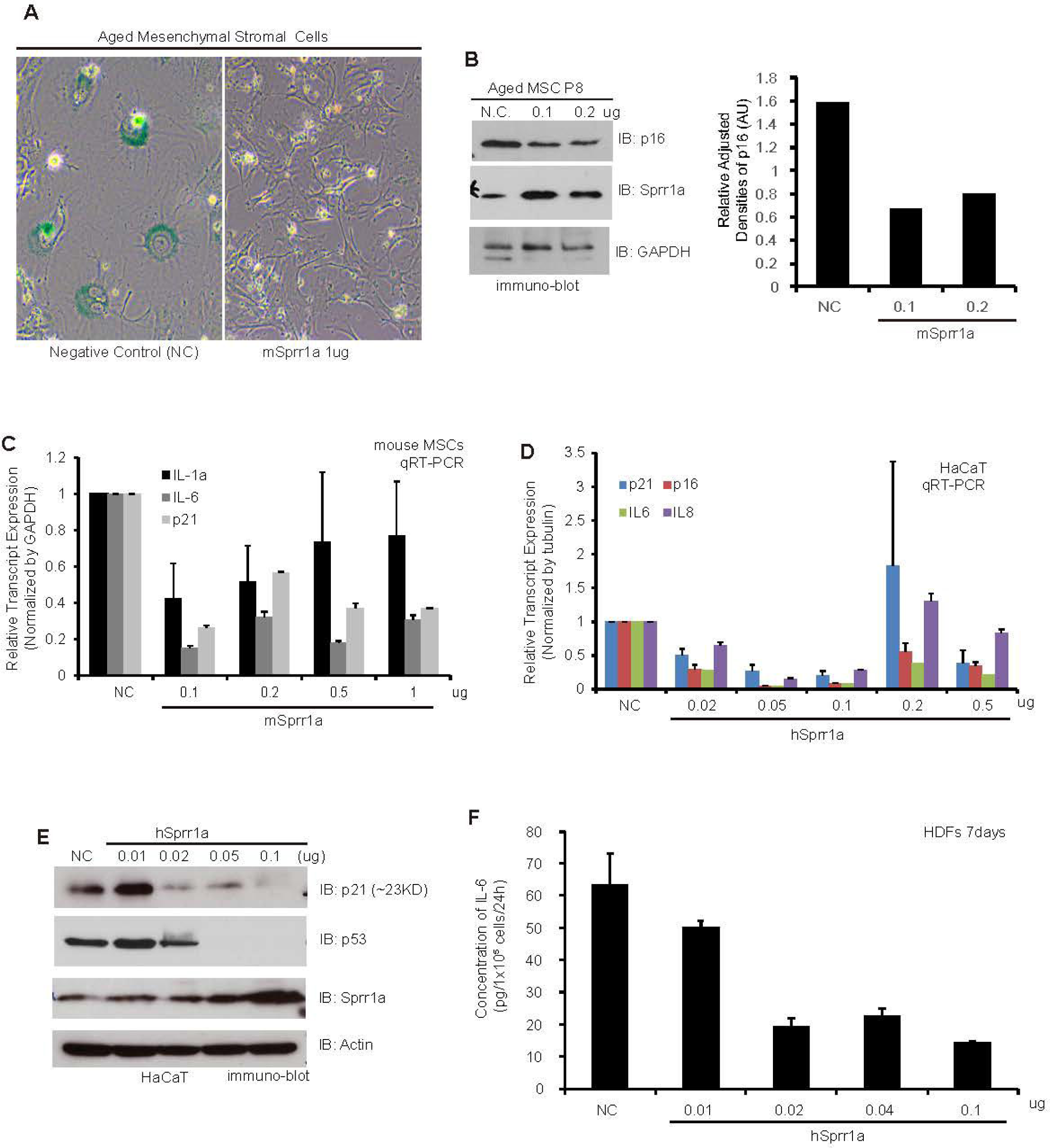
Sprr1a expression inhibits senescence-associated phenotypes. (A) Mouse Sprr1a was transiently expressed in mouse MSCs isolated from aged mice (18-22 months old). After transfection, cell shapes were carefully monitored and senescence evaluated *via* SA-β-galactosidase staining. (B) Following exogenous expression of Sprr1a (non-transfected versus 0.1 or 0.2ug), a major factor promoting senescence, p16, was evaluated *via* immuno-blotting. GAPDH was employed as an internal control. Densitometry (right panel) shows p16 levels normalized to actin. N.C. is transfected empty control vector. (C) Following Sprr1a expression at the indicated levels in mouse MSCs, the transcript levels of another cell cycle regulator, p21, as well as SASP factors, IL-6 and IL-1a, were monitored using real-time PCR. (D) To evaluate the role of Sprr1a in human cells, human Sprr1a was transiently expressed in HaCaT. The transcript levels of p16, p21, IL-6 and IL-8 were observed using real-time PCR. (E) After the transfection of Sprr1a at the indicated levels, the protein levels of p21 and p53 were evaluated using immuno-blotting. Actin is employed as a loading control. (F) Human Sprr1a was exogenously expressed in HDF cells. Cells were cultured for 7days after transfection to study the senescent phenotype. Conditioned media (CM) from cells transfected with the indicated levels of Sprr1a were analyzed for IL-6 by ELISA. The concentration of IL-6 was normalized by cell number. Shown is a representative example of two experiments. Error bars represent the standard deviation of triplicate determinations.

Primary mouse MSCs grow relatively slowly. To help circumvent this issue in examining Sprr1a, as well as to utilize independent cell and species sources, we tested the human HaCaT cell line and primary human dermal fibroblasts (HDFs). While human Sprra1 is smaller than mouse Sprra1, of those residues that human Sprra1 contains, 87% are identical between human and mouse (TCoffee and NIH BLAST) (Supplemental Fig 1B). As seen in mouse MSCs, p21 transcript levels were decreased upon the expression of human Sprr1a in HaCaT cells, and this was likewise observed for transcript levels of p16 RNA (Fig 1D) (not statistically significant was an unexpected increased of p21 seen at 0.2 ug hSprr1a). The transcript levels of IL-6 and IL-8, which are SASP factors, additionally became reduced upon Sprr1 expression in HaCaT cells (Fig 1D). For unknown reasons, such effects upon IL-8 in particular were not apparent at higher transfection levels of hSprr1a (0.2ug and 0.5ug). When evaluated at the protein level, p21 and p53 were graphically lowered in a dose-dependent manner upon Sprr1a’s expression (Fig 1E). We also evaluated IL-6 secreted from human dermal fibroblasts (HDFs) made to express Sprr1a. Since IL-6 levels can be detected 6 or 7 days after treatment with senescence-inducing factors [41], ELISA was used to monitor IL-6 levels in the medium 7 days post transfection with Sprr1a. As expected, Sprr1a expression resulted in the dose-dependent lowering of IL-6 levels (Fig 1F). These data suggest that Sprr1a can reverse markers of cellular senescence, and by extension, potentially contribute to rejuvenation.

### Senescence-like phenotype is promoted via the depletion of Sprr1a

We next tested whether the knockdown of Sprr1a expression induces senescence-like phenotype. Each of three independent siRNAs directed against Sprr1a reduced its mRNA levels (Fig 2A). Among them, siSprr1a-3 showed a marked decrease, so we selected this siRNA to test further. Complementing what was seen upon Sprr1a expression (Fig. 1), the depletion of Sprr1a instead increased IL-6 protein levels in human dermal fibroblasts (HDFs) (Fig 2B). In response to various stimuli including replicative exhaustion and radiation, cellular senescence is a process that imposes permanent proliferative arrest. Thus, we examined the effect of Sprr1a knockdown on cell proliferation. Using the IncuCyte cell count proliferation assay, we found that human dermal fibroblasts depleted of Sprr1a exhibit decreased levels of proliferation compared to controls (Fig 2C). Consistent with our proliferation assays, Sprr1a-depleted human dermal fibroblasts evidenced greater SA-β-galactosidase staining relative to control siRNA transfected cells (Fig 2D). We then assessed the distribution of intracellular HMGB1 (high mobility group box 1), as it is reported to relocalize outside the nucleus in senescent human and mouse cells in culture and *in vivo* [42]. Correspondingly, we found that there is an increased appearance of HMGB1 in the cytoplasm upon siRNA-mediated depletion of Sprr1a (Fig 2E). In addition to employing siRNA (above), we tested the impact of shRNA-mediated knock-down of Sprr1a in mouse MSCs. Following viral driven delivery of the shRNA, MSCs displayed the anticipated senescence-like phenotype of increased cell spreading relative to controls (Supplemental Fig 2A and 2B). Such findings are consistent with the promotion by Sprr1a of non-senescent/ young phenotypes.

**Figure 2.**
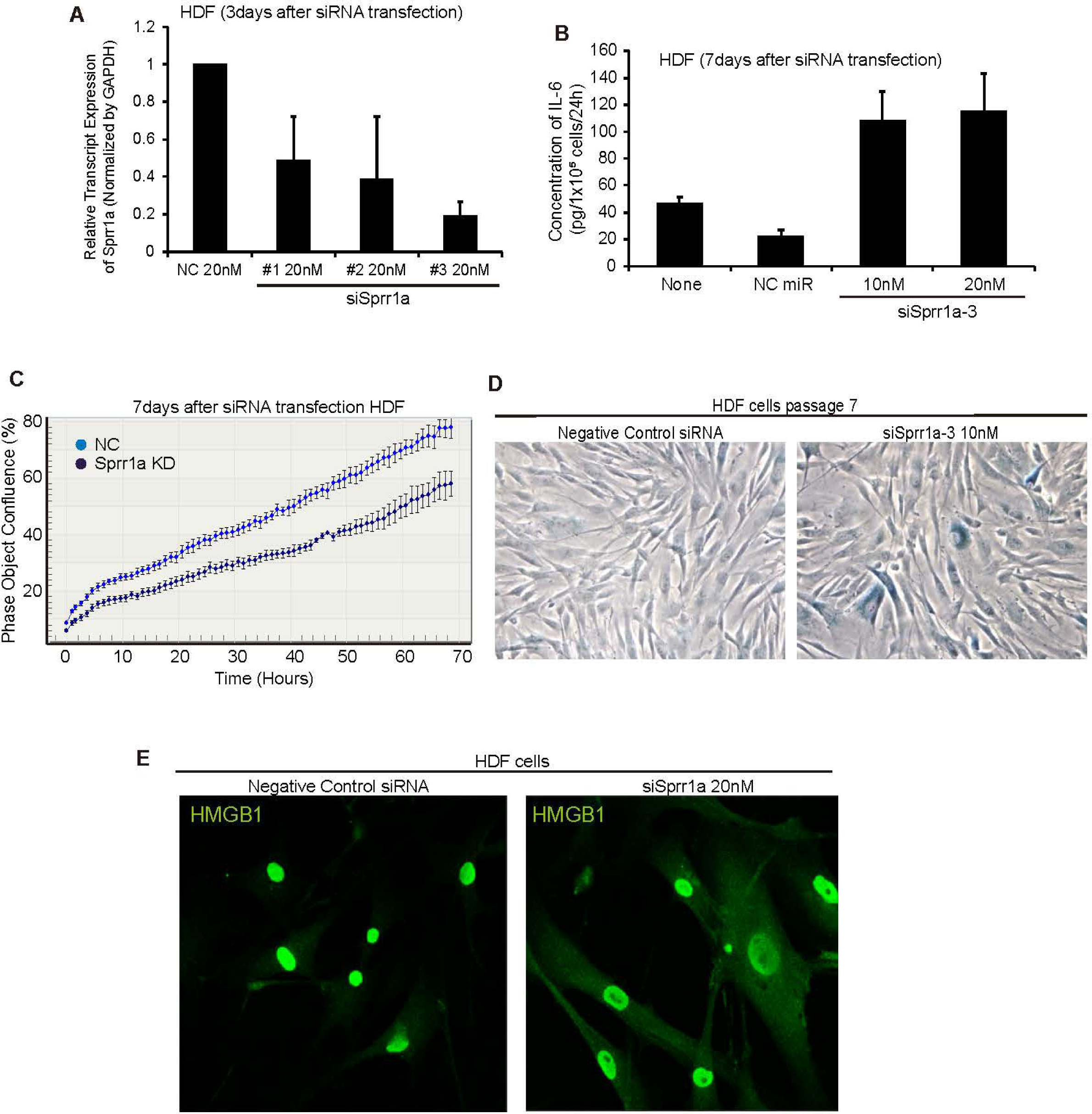
Cellular senescence is promoted via the depletion of Sprr1a. (A) Lysates from HDF cells transfected with three distinct Sprr1a siRNAs were analyzed for Sprr1a using Real-time PCR. GAPDH was used as a negative control. (B) The indicated amounts of human Sprr1a were transfected into HDF cells, that were then cultured for 7 days to study the senescent phenotype. Conditioned media (CM) was collected from HDF cells and used for ELISA to detect IL6’s level. NC means negative control siRNA (Qiagen, SI03650318). Shown is a representative of two experiments. (C) IncuCyte cell confluence proliferation assay was performed with HDF cells, in which 10nM of NC or siSprr1a is transfected. Cells were seeded into 96-well plate after transfection and images captured every 1 to 2 hours using an IncyCyte ZOOM system. (D) Senescent phenotypes 7 days following the transfection of siSprr1a were monitored using β-galactosidase assays. (E) Sprr1a siRNA was transfected into HDF cells, later fixed with 4% PFA for 10 minutes, blocked with BSA in PBS and immuno-stained with antibody against HMGB1. Each experiment was repeated three or more times. Error bars represent the standard deviation of triplicate determinations.

### miR-130b modulates the expression of Sprr1a to regulate cell senescence

Following the use of microarrays to point to microRNAs whose levels vary significantly with aging, we employed *in silico* analyses (MicroCosm) to suggest those that might target transcripts of Sprr1a. From the arrays, miR-130b-3p is expressed at approximately 2-fold greater levels in old *versus* young MSCs, and based upon their respective sequences, mouse miR-130b-3p further appears to have the potential to target a region within the 3’ sUTR of mouse Sprr1a. While miR-130b has been studied in relation to cancer, there is not yet agreement as to its role [43–47]. One interesting report indicates that the miR-130b∼301 cluster activates cell cycle inhibitors epigenetically, followed by SASP [48]. In addition, miR-130b expression is observed to increase during replicative senescence [49]. To further assess our microarray data, we used qRT-PCR to look at the level of miR-130b-3p in young *versus* aged MSCs. While subtle, we confirmed slightly elevated levels of miR-130b-3p in aged mouse MSCs (Fig 3A). This trend was observed for MSCs isolated independently from each of two mice. We next evaluated if miR-130b-3p and/ or miR-130b-5p modulate Sprr1a levels. The nucleotide sequences of miR-130b-3p and −5p complement each other, being derived from the right *versus* left arms of the pre-microRNA-130b (Supplemental Fig 1C). Our initial expectation was that −5p might serve as a negative control for the activity of −3p. Measured relative to the effect of an accepted standard miR negative control (NC) (Qiagen catalog # 1022076), and normalized to the actin loading control, our immuno-blots indicated that the expression of mouse miR-130b-3p (less so mouse −5p) led to ∼10-fold decreases in Sprr1a protein levels in mouse MSCs (Fig 3B). We also evaluated Sprr1a at the mRNA level and found that the exogenous expression of miR-130b-3p decreased Sprr1a up to ∼20-fold in mouse MSCs (Fig 3C). Unexpectedly, mouse miR-130b-5p also exhibited notable effects upon Sprr1a’s RNA levels in mouse MSCs (Fig 3C), with a lesser effect being observed upon Sprr1a’s protein levels (Fig. 3B). Further, we noted that human miR-130b-5p exhibited an impact upon the mRNA levels of Sprr1a when expressed in human primary dermal fibroblasts (HDFs) (Fig 3D), and that mouse miR-130b-5p in common with mouse miR-130b-3p induced cellular senescence as measured by β-galactosidase staining in mouse MSCs at 6-7 days (Fig 3E). In general, mouse miR-130b-3p appeared to have a greater impact upon Sprr1a’s protein levels in mouse MSCs, whereas human miR-130b-5p appeared more effective in HDFs (comparing Fig 3B, 3C and 3D). The observation that both miR-130b-5p and −3p exhibit activity might reflect recent findings that both arms of pre-miRNAs are capable of generating mature miRNAs. For example, it is referred to as “arm switching” when each of the two arms is observed to generate an effect according to the context, such as in relation to the species or developmental stage [50–52]. In either case, miR-130b-5p and −3p each had an effect upon Sprr1a levels relative to the standard negative-control miRNA (NC), as well as to additional tested negative-control miRNAs (e.g. miR-22-3p in Fig. 3F). Since p53 and p16 protein levels are increased following the exogenous expression of miR-130b-3p or −5p (Fig 3F), similar to the depletion of Sprr1a (data not shown), our findings are again consistent with the possibility that miR-130b modulates senescence-like phenotype *via* Sprr1a. Further, in common with Sprr1a depletion, miR-130b-5p or −3p expression promotes the appearance of β-galactosidase staining in HDFs (Fig 3G), suggesting that miR-130b is related to senescence. In further keeping with this possibility, exogenous expression of miR-130b-3p and −5p in HDFs reduced the proportion of cells in S phase (Supplemental Fig 3A), and the exogenous expression of miR-130b increased trans-localization of HMGB1 (Supplemental Fig 3B), similar to what was seen earlier upon the knock-down of Sprr1a (Fig 2E).

**Figure 3.**
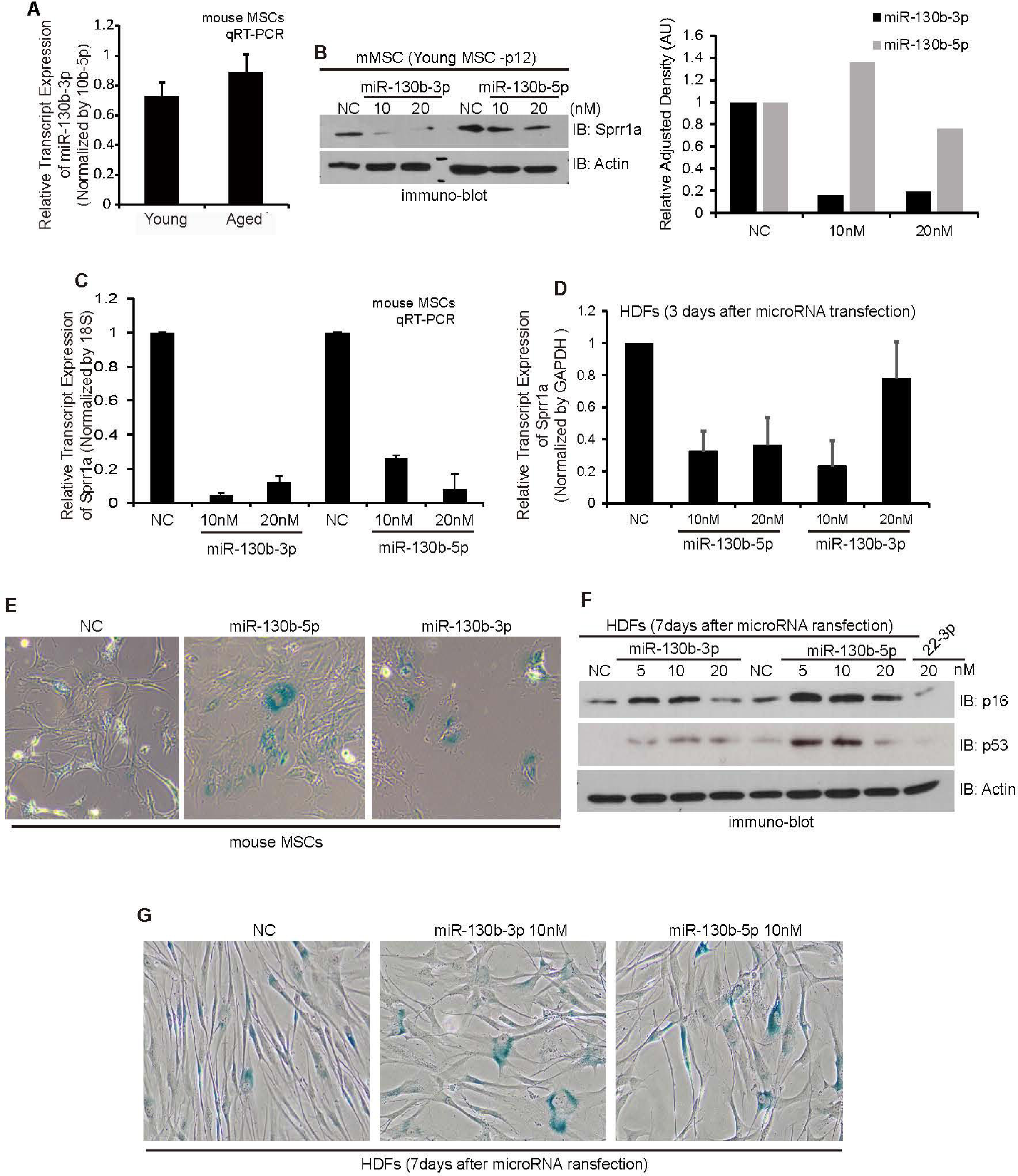
Mir-130b modulates the expression of Sprr1a and cellular senescence. (A) Young and aged MSCs, isolated from young (2-3 months) versus aged mice (18-20 months), were cultured *in vitro*. Mouse MSCs (mMSCs) were harvested and cDNA was subject to RT-PCR to assay mouse miR-130b-3p levels. (B) Endogenous protein levels of Sprr1a was monitored *via* immuno-blot (IB) in response to mouse miR-130b-3p or miR-130b-5p. Actin serves as loading control. Right panel: employing densitometry and ImageJ software, endogenous Sprr1a levels were quantitated and normalized relative to actin loading control. (C) The indicated amounts of mouse miR-130b-3p or miR-130b-5p were transfected into mouse MSCs. The corresponding cDNA was isolated to perform quantitative RT-PCR to monitor Sprr1a levels. (D) HDF cells were transfected with human miR-130b-3p or −5p for 48 hours. Endogenous Sprr1a levels were monitored through qRT-PCR. (E) Mouse MSCs were transfected with miR-130b-5p or 3p and the cells subjected to β-galactosidase assays. (F) HDF cells were transiently transfected with human miR-130b-5p or −3p. The effects were assessed by means of p16, p53 and actin immuno-blotting 7 days after transfection. (G) Senescence associated β-gal staining is enhanced upon the ectopic expression of human miR-130b-5p or −5p in HDF cells. Error bars here represent the standard deviation of triplicate determinations.

### Antagomirs of miR-130b inhibit the capacity of miR-130b to promote senescence-like phenotype

Based upon the senescence promoting properties of miR-130b-5p and −3p, exogenous expression of the corresponding antagomirs would be expected to inhibit senescence-like phenotype. Indeed, use of antagomirs to miR-130b resulted in increased levels of Sprr1a, implying again that Sprr1a is functionally linked to miR-130b (Fig 4A). Complementing our previous observations made in mouse MSCs, an antagomir directed against miR-130b-3p is more effective than an antagomir against −5p in raising the levels of Sprr1a (Fig 4A), with the converse being observed in HDF cells (data not shown). We next tested the protein levels of p16. As predicted using HDF cells (antagomirs against −5p being more effective than those against −3p in protecting Sprr1a), p16 levels were partially lowered (Fig 4B). Further, when we limited senescence in HDF cells by expressing antagomirs against miR-130b, we observed a reduction of the pro-senescence cytokine IL-6 (Fig 4C).

**Figure 4.**
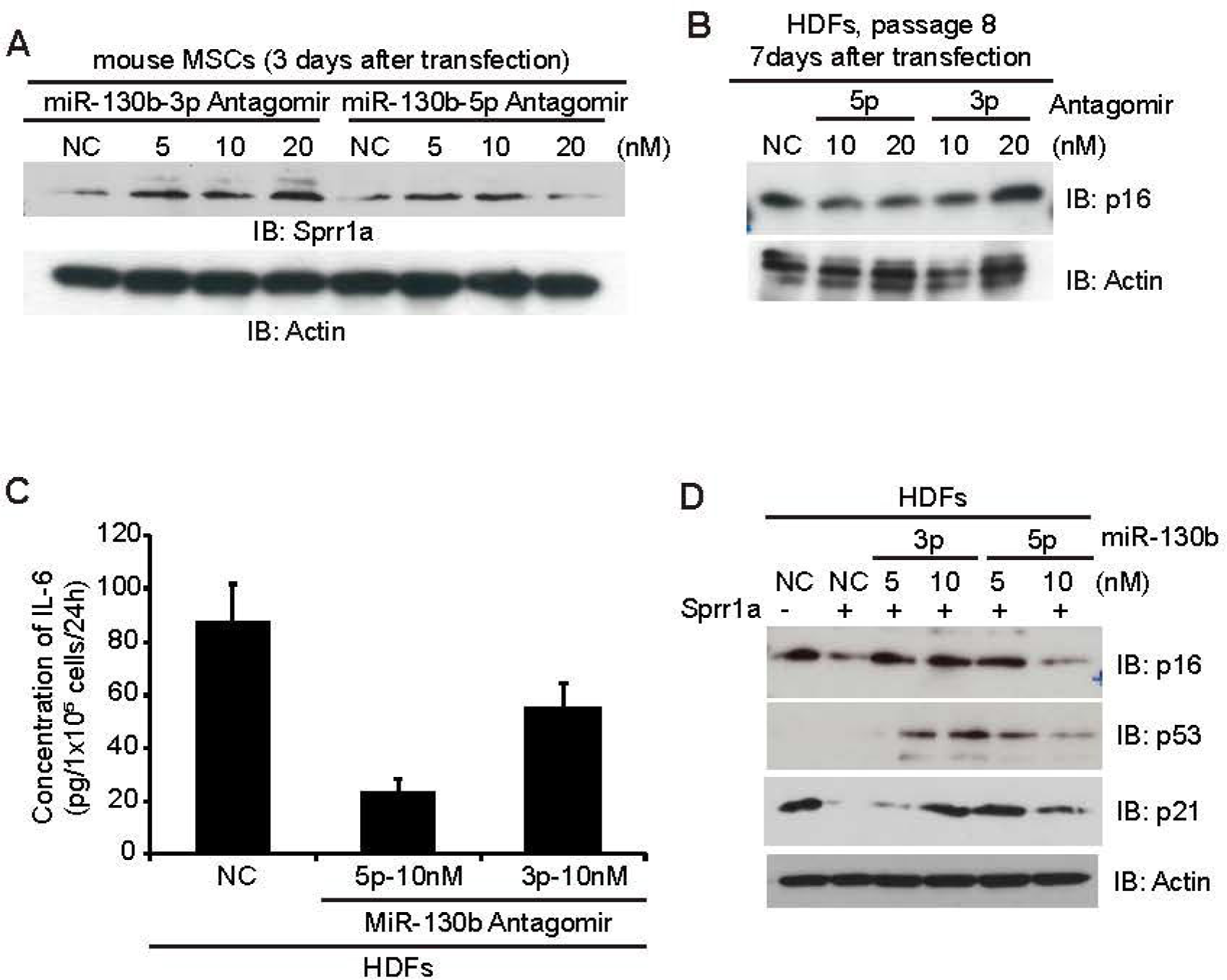
Antagomirs of miR130b inhibit its capacity to promote senescence. (A) Mouse MSCs were transfected with mouse antagomirs directed against miR-130b-3p or −5p. Three days after transfection, cells were harvested and endogenous Sprr1a levels were monitored *via* immuno-blotting (IB). Actin was employed as a loading control. (B) The indicated amounts of antagomirs were ectopically expressed and the HDF cells maintained for 7days. Cells were harvested and p16’s level observed relative to actin, a negative control. (C) After transfecting the antagomirs, HDF cells were cultured for 6 days. One day prior to cell harvest, low serum medium (1% FBS) was provided for 24 hours. The medium was then collected and assayed for IL-6 levels using ELISA (R&D systems). Error bars represent the standard deviation of triplicate determinations. (D) Human Sprr1a was transiently expressed into HDF cells with microRNA mimics as indicated. Cells were harvested 3 days after transfection and the expression levels of p16, p53 and p21 were monitored using immuno-blotting (IB). Actin served as a loading control.

Also at the functional level, we went on to test if miR130b-3p and −5p were capable of interfering with Sprr1a’s ability to counter senescence-like phenotype. Indeed, the reduced levels of p16, p53 and p21 following Sprr1a expression (as observed earlier), become increased (rescued) upon the co-expression of miR-130b (Fig 4D). Together with our earlier findings, such a rescue by miR-130b suggests that it may modulate the senescence-like phenotype *via* Sprr1a. Since the Sprr1a construct that we expressed contains the coding region but not the Sprr1a 3’ and 5’ UTRs, we anticipated being able to use it to rescue the senescence-like phenotype induced by miR130. However, since such rescue failed, we expect that the observed regulation of Sprr1a by miR-130b may be indirect.

We further observed that endogenous Sprr1a is present in both the nucleus and cytoplasm, whereas exogenous Sprr1a is largely localized to the cytoplasm (data not shown). Conceivably, endogenous nuclear Sprr1a might have roles in senescence-like phenotype that are unrelated to miR-130b. Future study will be needed of the detailed mechanisms regulating Sprr1a in cellular senescence.

### Senescence-like phenotype is inhibited upon exogenous expression of Sprr1a

To examine whether Sprr1a can reverse or limit cellular senescence-like phenotype, we first induced HDF cells to senesce using a standard program of exposure to X-irradiation. This stimulus caused a striking loss of Sprr1a levels, as followed using real-time PCR (Fig 5A). To examine the effect of Sprr1a on induced senescence-like phenotype, Sprr1a was ectopically expressed in primary HDFs followed by X-irradiation. As seen in other reports [53], X-irradiation increased p16’s protein levels. As we predicted, in a dose-dependent manner, Sprr1a expression limited the increase in X-irradiation-induced p16 levels (Fig 5B). We tested other stimuli of cellular senescence such as Doxorubicin. Primary HDFs were exposed to Doxorubicin for 4 days and real-time PCR performed to monitor Sprr1a. Sprr1a levels decreased in a dose-dependent manner (Fig 5C), implying that Sprr1a levels are lowered upon exposure to senescence stimuli. We also tested whether the expression of Sprr1a can inhibit replicative senescence-like phenotype (compare the upper left two images in Fig 5D). In this experiment, we employed HDFs at passage 11, which were then allowed to undergo a number of further passages to induce replicative senescence. Thus, the negative control (NC) cells exhibited strong β-gal staining. Simultaneously, we checked whether the expression of Sprr1a counters senescence promoted by cell incubation with Doxorubicin (compare upper and lower images in Fig 5D). As expected, exogenous Sprr1a lowered the SA-β-gal staining otherwise promoted by replicative senescence or by Doxorubicin, implying that Sprr1a may reduce senescence-like phenotype (Fig 5D & E).

**Figure 5.**
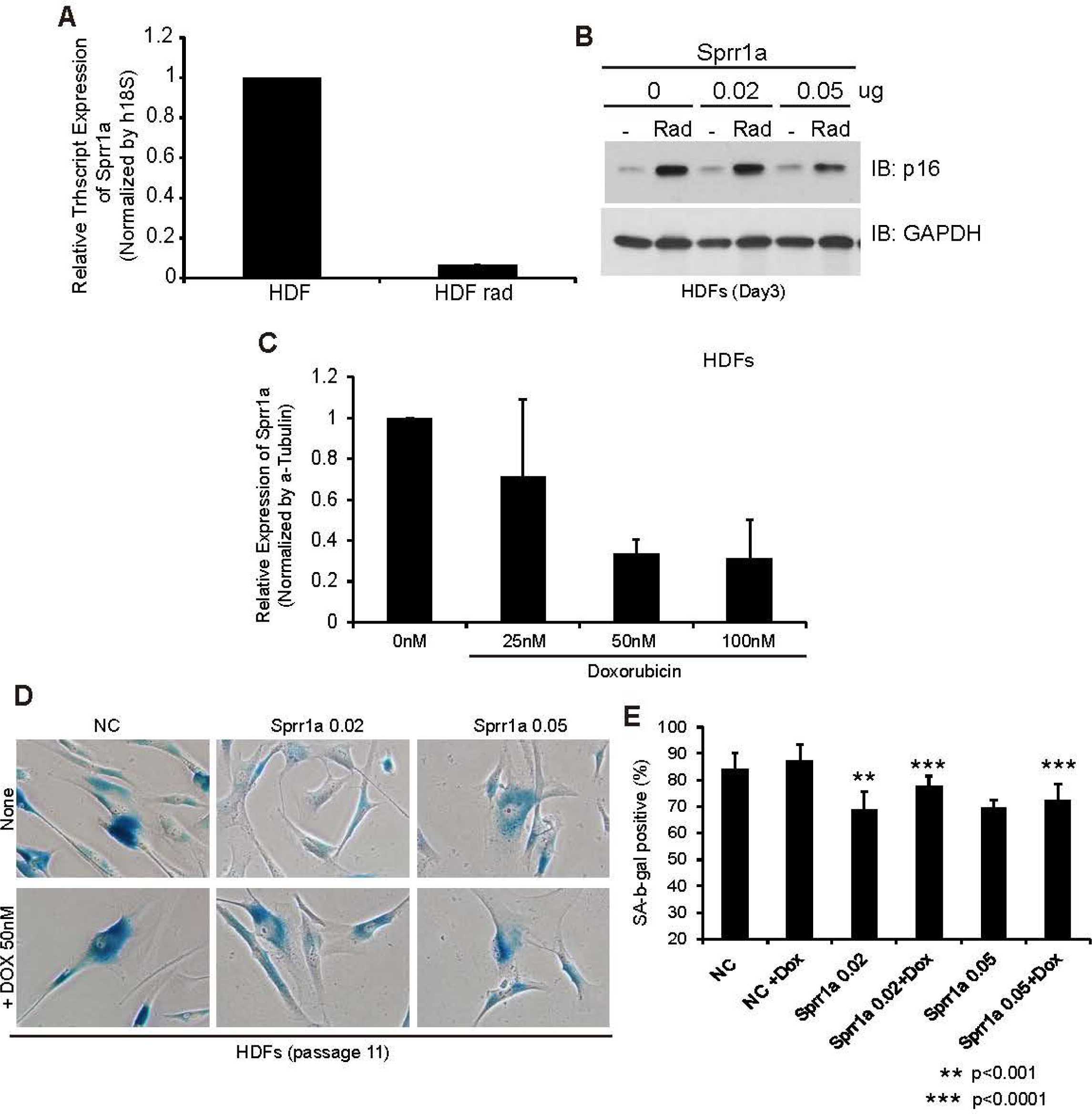
Senescence is inhibited upon exogenous expression of Sprr1a. (A) Human dermal fibroblasts (HDF) were irradiated at 10Gy. RNA was extracted 3 days following irradiation and Sprr1a levels assessed *via* quantitative real-time PCR. (B) Human Sprr1a was transiently expressed in HDF cells and cultured for 2 days. On the third day, cells were radiated at 10 Gy. Cells were harvested on the fifth day to observe p16 levels *via* immuno-blotting (IB). (C) Using HDF cells, Doxorubicin was present in the cell medium over four days. RNA was isolated and the transcript level of Sprr1a was monitored via real-time PCR. Error bars represent the standard deviation of triplicate determinations. (D & E) Sprr1a was transiently expressed in HDF cells and cells were treated with Doxorubicin three days after transfection. Doxorubicin was present in the cell medium over two days. Cells stained with β-gal were counted. Error bars represent the standard deviation of triplicate determinations. Student’s t-test was used to compare two groups and P<0.05 was regarded as statistically significant.

## DISCUSSION

To the best of our knowledge, we here report the first demonstration of Sprr1a’s role in senescence-like phenotype in cells. The ectopic expression of Sprr1a changed the morphology of aged MSCs to more resemble that of younger MSCs. Cell cycle regulators and SASP factors decreased when Sprr1a was overexpressed. Employing human primary cells such as HDFs we found that Sprr1a modulates aging-related factors, reflected for example, in changes in the levels of p16 and p53 levels, cell proliferation and SA-β-galactosidase staining. Furthermore, our data suggest that miR-130b is as upstream indirect modulator of Sprr1a. For example, MiR-130b expression promoted cellular senescence-like phenotype and could rescue phenotypes induced by the exogenous expression of Sprr1a. We also revealed that Sprr1a-expressing cells show less response to Radiation and Doxorubicin, two agents otherwise known to promote cellular senescence. Taken together, we propose the existence of a new mir-130b/ Sprr1a pathway (or relationship), that is likely to contribute to senescence-like processes.

The mechanisms underlying how miR-130b regulates Sprr1a needs further evaluation. While expression of the Sprr1a coding region (lacks 5’ and 3’UTRs) failed to rescue phenotypic changes brought about by exogenous expression of miR-130b (data not shown), miR-130b was capable of rescuing the phenotype resulting from the expression of Sprr1a. This suggests that miR-130b potentially decreases the level of Sprr1a via an indirect mechanism. However, it is still possible that a miR130b target sequence resides somewhere within the coding region of Sprr1a.

Elucidation of the actual mechanisms accounting for Sprr1a’s downstream effects upon cellular senescence-like phenotype remains a task for future studies, as does defining Sprr1a’s upstream mechanistic links to miR-130b-3p and −5p. One published study upon cardiomyocytes suggests that Sprr1a is a downstream mediator of gp130 cytokines, and that Sprr1a expression provides a protective effect in response to ischemia-induced cardiac injury, possibly through stabilization of the actin cytoskeleton [39, 54]. As mentioned above, Sprr1a protein has also been named Cornifin-A, characterized as a cross-linked envelope protein of keratinocytes that first appears in the cell cytosol and becomes cross-linked by transglutaminase to the intracellular domains of membrane proteins associated with the plasma membrane. Since we find that expression of SPRR1A in MSCs alters the cells’ shape, coming to resemble that of young MSCs, SPRR1a is likely to modulate cell shape via an impact upon cytoskeletal structures relevant to aging phenotypes. While speculative, Sprr1a might for example participate in modulating or executing cell-shape responses that oppose those of senescence stimulators.

In addition, strongly increased expression of Sprr1a has been observed upon peripheral nerve injury [1] . Given that Sprr1a is a regeneration-associated protein, Sprr1a may suppress senescence-associated phenotypes via effects upon the cell cycle. Our evidence indicates that depletion of Sprr1a has a partial if not complete negative impact upon cell proliferation.

Alternatively, recent reports in mice indicate that inhibition of the mTOR pathway (w/ rapamycin) significantly extends lifespan and alleviates age-related disorders [41]. However, the connection between mTOR signaling and Sprr1a is unclear. There are several potential protein interactions of Sprr1a listed in open databases (e.g. BioGRID, InAct, STRING). However, none have been functionally linked to Sprr1a, with the exception of EGF, where Sprr1a levels increase upon treatment [55]. Thus, while future studies are needed to define the larger upstream and downstream pathways involved, our contribution here is in reporting the modulation of cell senescence by a newly revealed miR-130b ➔ Sprr1a regulatory relationship.

## METHODS

### Isolation of Bone marrow stromal cells

C57BL/6J mice were used to isolate bone marrow stromal cells. We cut the end of the tibia and femur just below the end of the marrow cavity into the spongy bone exposed by removal of the growth plate. The marrow was plugged out of the cut end of the bone with 1ml of complete media and Mesenchymal stromal cells (MSCs) were obtained from murine bone marrow by serial passage of adherent cells in Dubecco’s modified Eagle’s medium (Hyclone, SH30022.01) containing 10% FBS as described [56]. Employing FACS Aria II, cells were flow sorted (FACS) to obtain the CD45-CD31-Ter119-population. Anti-mouse CD45 (eBioscience, 11-0451), Anti-mouse CD31 (eBioscience, 11-0311), and anti-mouse Ter119 (eBioscience, 11-5921) antibodies were employed to label the cells.

### Mammalian cell culture

Human dermal fibroblasts (HDFs) were purchased from ATCC, authenticated as mycoplasma-free and maintained in Dulbecco’s modified Eagle’s medium containing 10% feral bovine serum.

### Transfection of plasmids, microRNAs and RNA interference

Using Lipofectamine 2000 (ThermoFisher, P/N 52887) or HDF transfection Avalanche transfection reagent (EZ biosystems, EZT-HDF0-1), mouse MSCs, HaCaT cells and HDF cells were transiently transfected with plasmid, microRNAs or siRNAs as described. After transfection, cells were lysed using RIPA buffer (ThermoFisher, #89900) at the indicated times, inclusive of a proteinase inhibitor cocktail (Roche, 11836170001). Constructs including human (NM_005987) and mouse Sprr1a (NM_009264) were purchased from Origene, bearing the pCMV6 vector backbone with C-terminal Myc-DDK Tag. Mimics of mmu-miR-130b-3p (Qiagen, MSY0000387), mmu-miR-130b-5p (Qiagen, MSY0004583), has-miR-130b-3p (Qiagen, MIMAT0000691) and has-miR-130b-5p (Qiagen, MIMAT0004680) were transiently expressed to induce senescence-like phenotype. Anti-has-miR-130b-3p (Qiagen, MIMAT0000691) and Ant-has-miR-130b-5p (Qiagen, MIMAT0004680) were employed to inhibit the respective microRNAs. Three different siRNAs of Sprr1a were purchased from IDT (NM-005987).

### Immunofluorescence staining

HDF cells transfected with the designated plasmid or microRNA were fixed with 4% paraformaldehyde (T&I, BPP-9004) and permeabilized with 0.3% Triton X-100 (Sigma-Aldrich, X100). Blocking was performed with 1% BSA (AMRESCO, 0332). Primary and secondary antibodies are employed to detect the indicated proteins. Cells on coverslip were stained with DAPI for nuclear staining. Confocal laser scanning images were acquired using a Leica 20X/0.7 NA objective lens and an DMI 6000 inverted microscope equipped with the Leica TCS SP8 STED CW system.

### SA-β-galactosidase staining

Senescence cells exhibit pH-dependent β-galactosidase activity. After the transfection of plasmid, siRNA or microRNA, cells were cultured for 7 days. A senescence β-galactosidase staining kit (Cell Signaling Technology, Cat# 9860) was employed to score for senescence. Cells were fixed and processed following the manufacturer’s instructions.

### ELISA assays

HDF cells were transfected with the indicated amounts of plasmids, microRNAs or siRNAs. Cells were cultured for 7 days to study the senescent phenotype, SASP. One day prior to cell harvest, low serum medium (1% FBS) was provided for 24 hours. Conditioned medium (CM) was collected from cells cultured serum-free for 24 hours, and was filtered and stored at - 80℃. We counted cell numbers for each sample and the concentration of IL-6 was normalized to cell numbers. An ELISA kit was used to measure IL-6 (R&D Systems) and we performed experiments following the manufacturer’s instructions.

### Real-Time PCR

Total RNA was prepared using Trizol (BIOLINE, BIO-38032). Tetro cDNA synthesis kit (BIOLINE, Cat# BIO-65043) was employed to synthesis cDNA. Primers for RT-PCR were: m18S, 5’-GTAACCCGTTGAACCCCATT-3’ and 5’-CCATCCAATCGGTAGTAGCG-3’; mGAPDH, 5’-TCACCACCATGGAGAAGGC-3’ and 5’-GCTAAGCAGTTGGTGGTGCA-3’; mIL-6, 5’-GCTACCAAACTGGATATAATCAGGA-3’ and 5’-CCAGGTAGCTATGGTACTCCAGAA-3’; mIL-1a, 5’-TTGGTTAAATGACCTGCAACA-3’ and 5’-GAGCGCTCACGAACAGTTG-3’; hIL-6, 5’-GCCCAGCTATGAACTCCTTCT-3’ and 5’-GAAGGCAGCAGGCAACAC-3’; human IL-8, 5’-AGACAGCAGAGCACACAAGC-3’ and 5’-ATGGTTCCTTCCGGTGGT-3’; human p16, 5’-GACCTGGCTGAGGAGCTG-3’ and 5’-GCATGGTTACTGCCTCTGGT-3’, ; p21, 5’-CAGAGGAGGCGCCAAGACAG-3’ and 5’-CCTGACGGCGGAAAACGC-3’; human GAPDH, 5’-GGTGGTCTCCTCTGACTTCAACA-3’ and 5’-GTGGTCGTTGAGGGCAATG-3’; human Tubulin, 5’-CTTCGTCTCCGCCATCAG-3’ and 5’-TTGCCAATCTGGACACCA-3’; human 18S rRNA, 5’-CTACCACATCCAAGGAAGCA-3’ and 5’-TTTTTCGTCACTACCTCCCCG-3’. To confirm the expression of miR-130b-p5 (Qiagen, MS00001554) and −3p (Qiagen, MS00011123), we purchased commercial primers from Qiagen. Primers for mouse Sprr1a (GeneCopoeia, MQP028278) and human Sprr1a (GeneCopoeia, HQP060361) were employed to detect expression of Sprr1a. For Real-time PCR, *Power* SYBER Green PCR Master Mix (ThermoFisher, 4367659) and GTaq® qPCR Master Mix (Promega, Cat.# A6001/2) were utilized.

### IncuCyte cell confluence proliferation assay

IncuCyte® ZOOM Live-Cell Analysis System (Essen Bioscience, Ann Arbor, Michigan) was employed to evaluate cell proliferation. HDF cells were transfected with the indicated siRNA or microRNA. After transfection, cells were cultured for three days, and 5,000 cells were seeded into 96-well plates provided from Essen Bioscience. Five hours after seeding, the proliferation of HDF cells was analyzed via Phase Object Confluence of the images, following instructions recommended by IncuCyte.

### Immunoblot analysis

Immuno-blotting employed commercial monoclonal antibodies directed against Sprr1a (ThermoFisher), p16 (Abcam), p53 (Santa Cruz, Cat# sc-6243), p21 (Santa Cruz, C-19), Actin (Santa Cruz), and GAPDH (Abcam). For immuno-blotting, cells were transfected with 0.5-2ug of the indicated plasmid DNA (10-15ug in the case of 100mm dishes), siRNA (5-20nM of 6well dishes) and/ or micro-RNA (5-20nM of 6well dishes). Mammalian cell lysates were prepared using RIPA buffer (1% NP40; ThermoFisher Scientific), inclusive of Halt^TM^ Protease and Phosphatase inhibitor cocktail (ThermoFisher Scientific). Cell equivalents were resolved by SDS-PAGE and transferred onto nitrocellulose membranes. Immunoblotting and antibody incubations took place in 2% bovine serum albumin-TBST (T&I). Super Signal West Pico reagents were utilized to detect HRP-conjugated secondary antibodies.

## CONFLICT OF INTEREST

The authors declare no competing financial interests.

## FUNDING

Our funding sources supporting this work are much appreciated: National Research Foundation of Korea, Ministry of Science, ICT & Future Planning NRF-2015R1C1A1A02036506 (JYH) and NIH/NIGMS 1 RO1 GM 107079-01A (PDM). Our DNA-sequencing was facilitated by an NIH/ NCI Core Grant (CA-16672) to UT MDACC.

## Notes

### Competing Interest Statement

The authors have declared no competing interest.

