## Supplemental Figure legends for "Sprr1 and miR-130b contribute to the senescence-like phenotype in aging"

**Supplementary Figures**

**Supplemental Figure 1. Phenotype of young versus aged mouse MSCs and a diagram of Sprr1a amino acid sequence alignment.** (A) Bone marrow from young (4-6 months) or aged (18-26 months) C57BL/6J mice was obtained to isolate bone marrow stromal cells. As described in M&M, mesenchymal stromal cells (MSCs) were obtained from murine bone marrow by serial passage of adherent cells in medium containing 10% FBS (HyClone, SH 30071.03). After the initial sub-culture, cells were monitored for aging phenotypes. (B) Sequence alignment of human and mouse Sprr1a. Identical and similar (conservative substitutions) residues are indicated with black highlighting. (C) Sequence alignment of human and mouse microRNA-130b. Left arms reflect miR-130b-5p and Right arms miR-130b-3p.

**Supplemental Figure 2. Sprr1a knock-down induces cellular senescence.** (A) Lentiviral transduction particles of sh-Sprr1a to knock down Sprr1a (Sigma, TRCN0000098406) were purchased from Sigma and used to infect mouse mesenchymal stromal cells. Cells were cultured in puromycin-containing medium for one week. Endogenous Sprr1a’s level were monitored *via* immuno-blotting (IB). Actin serves as a negative control. (B) Images of sh-empty versus sh-Sprr1a infected cells were taken after 1 week of puromycin selection. Yellow stars indicate senescence phenotypes resulted from the infection of sh-Sprr1a.

**Supplemental Figure 3. MiR-130b promotes senescence phenotype.** (A) To address the effect of miR-130b upon the cell cycle, mouse MSCs were transfected with miR-130b-3p mimics, cultured for 7 days, and subjected to flow cytometry employing the BD Pharmigen BrdU Flow Kit (Cat# 559619). In brief, 10ul BrdU was added directly to each mL of tissue culture medium and incubated. MSCs were fixed and permeabilized. Total DNA was stained with BrdU, and with 7-AAD. Cell cycle distributions were monitored using FACS LSRII. (B) MiR-130b-5p or -3p was transfected into HDFs. Cells were cultured for 4 days and the immunofluorescence localization of HMGB1 was monitored employing anti-HMGB1 antibody (Abcam, ab18256,).
