## Supplemental Figures for "Sprr1 and miR-130b contribute to the senescence-like phenotype in aging"

Supplement Figure 1 - Hong et al.
Phenotype of young versus aged mouse MSCs and
a diagram of Sprr1a amino acid sequence alignment

a

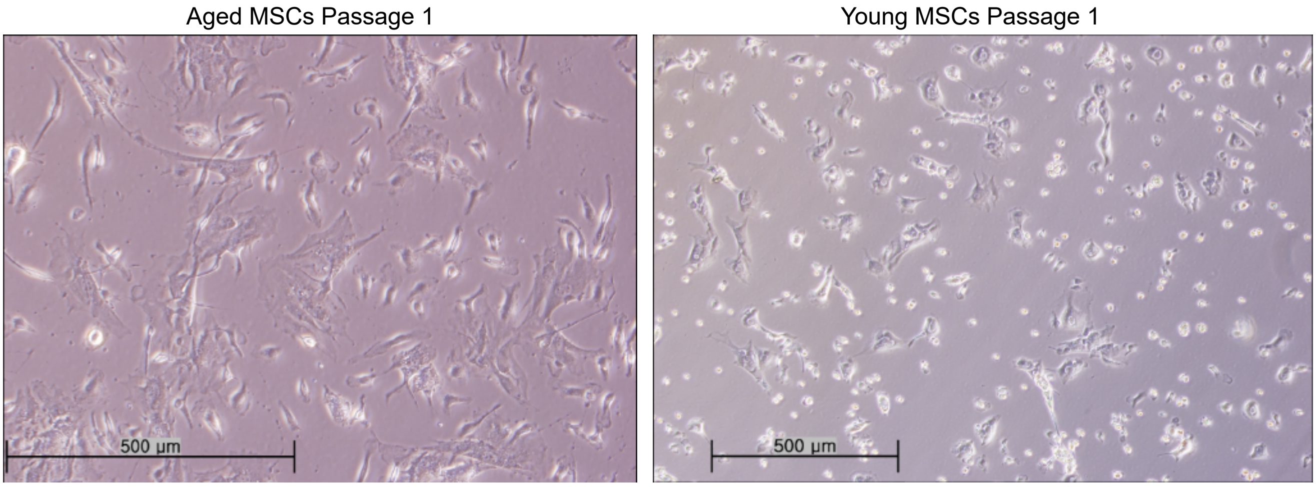


b

```
Human     1  MNSQQQKQPCTPPPQPQQQQVKQPCQPPPQEPCIPKTKEPCH------------------
Mouse     1  MSSHQQKQPCTVPPQLHQQQVKQPCQPPPQEPCAPKTKDPCHPVPEPCNPKGPEPCHPKA

Human    43  -----PKVPEPCHPKVPEPCQPKVPEPCQPK-----------------------------
Mouse    61  PEPCHPKAPEPCNPKVPEPCQPKVPEPCQPKVPEPCNPKVPEPCQPKAPEPCHPKAPEPC

Human    69  ---VPEPCPSTVTPAPAQQKTKQK
Mouse   121  HPVVPEPCPSTVTPSPYQQKTKQK
```

c

MiR-130b

```
Human  5′-GGCCUGCCCGACACUCUUUCCCUGUUGCACUACUAUAGGCCGCUGGGAAG
Mouse  5′-GGCUUGUUGGACACUCUUUCCCUGUUGCACUACUGUGGGCCUCUGGGAAG

Human      CAGUGCAAUGAUGAAAGGGCAUCGGUCAGGUC-3′
Mouse      CAGUGCAAUGAUGAAAGGGCAUCUGUCGGGCC-3′
```

Supplement Figure 2 - Hong et al.
Sprr1a knock-down induces cellular senescence

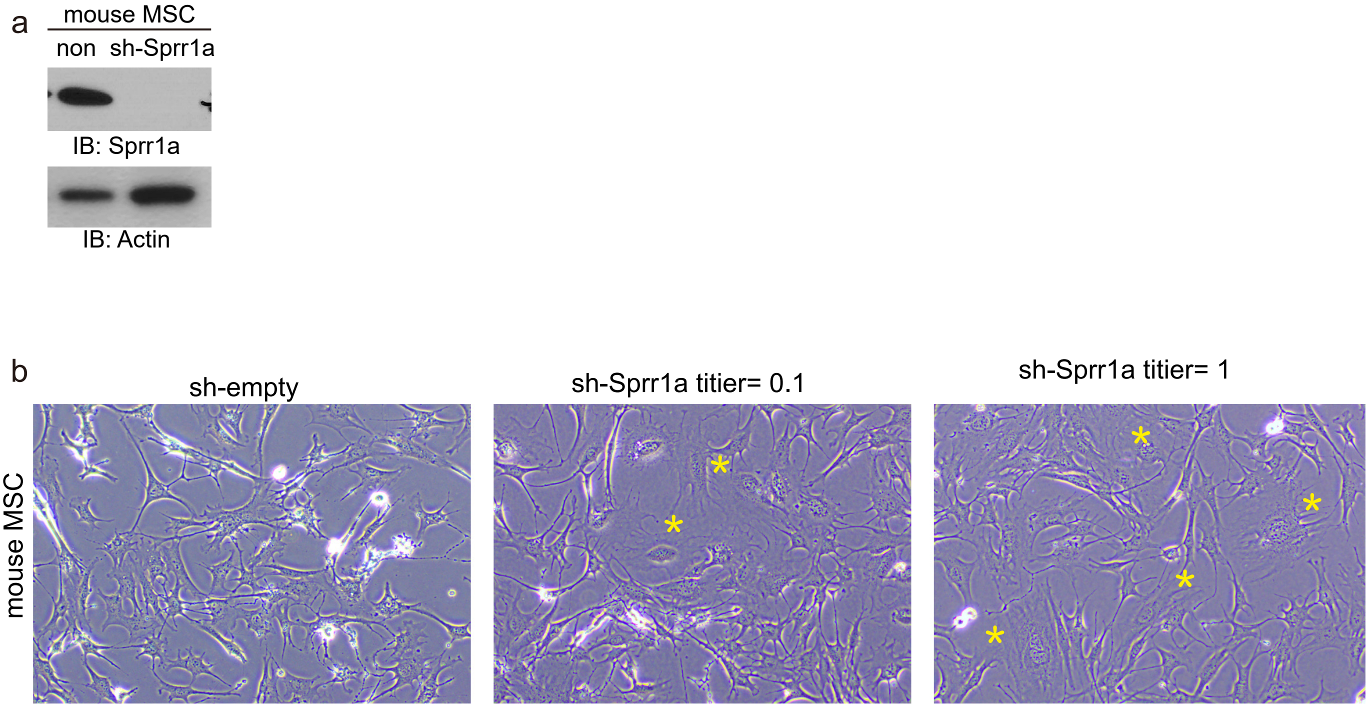

Supplement Figure 3 - Hong et al.
MiR-130b promotes senescence phenotype

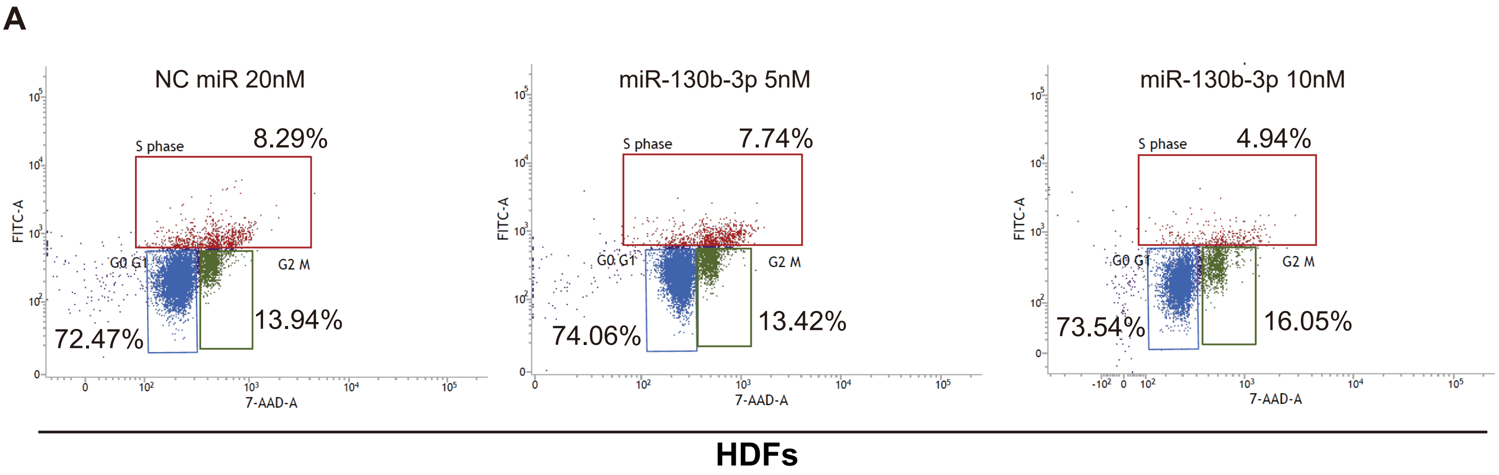


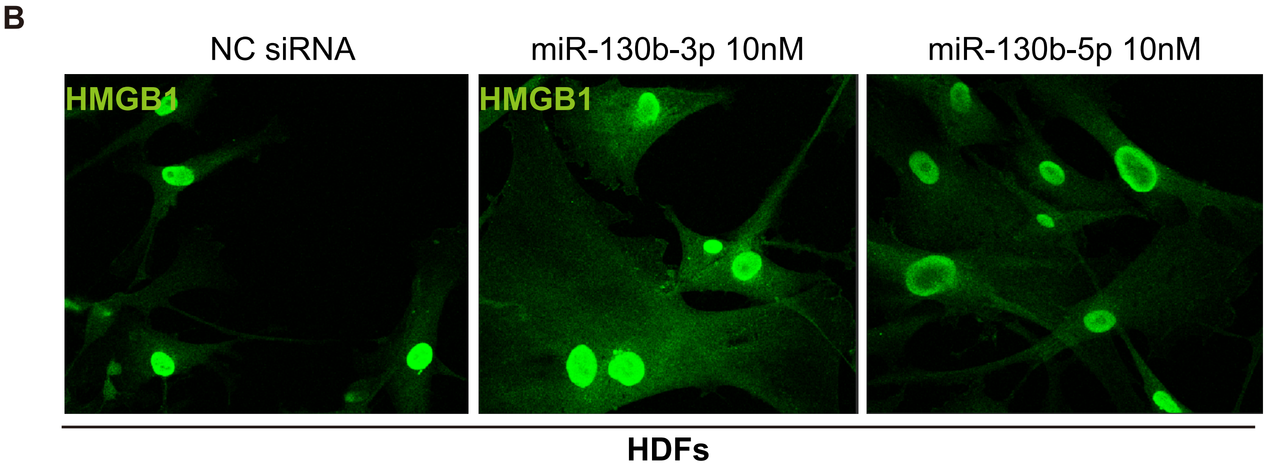
